## Supplemental Information for "Sequential action of a tRNA base editor in conversion of cytidine to pseudouridine"

^6^Singapore-MIT Alliance for Research and Technology Antimicrobial Resistance Interdisciplinary Research Group

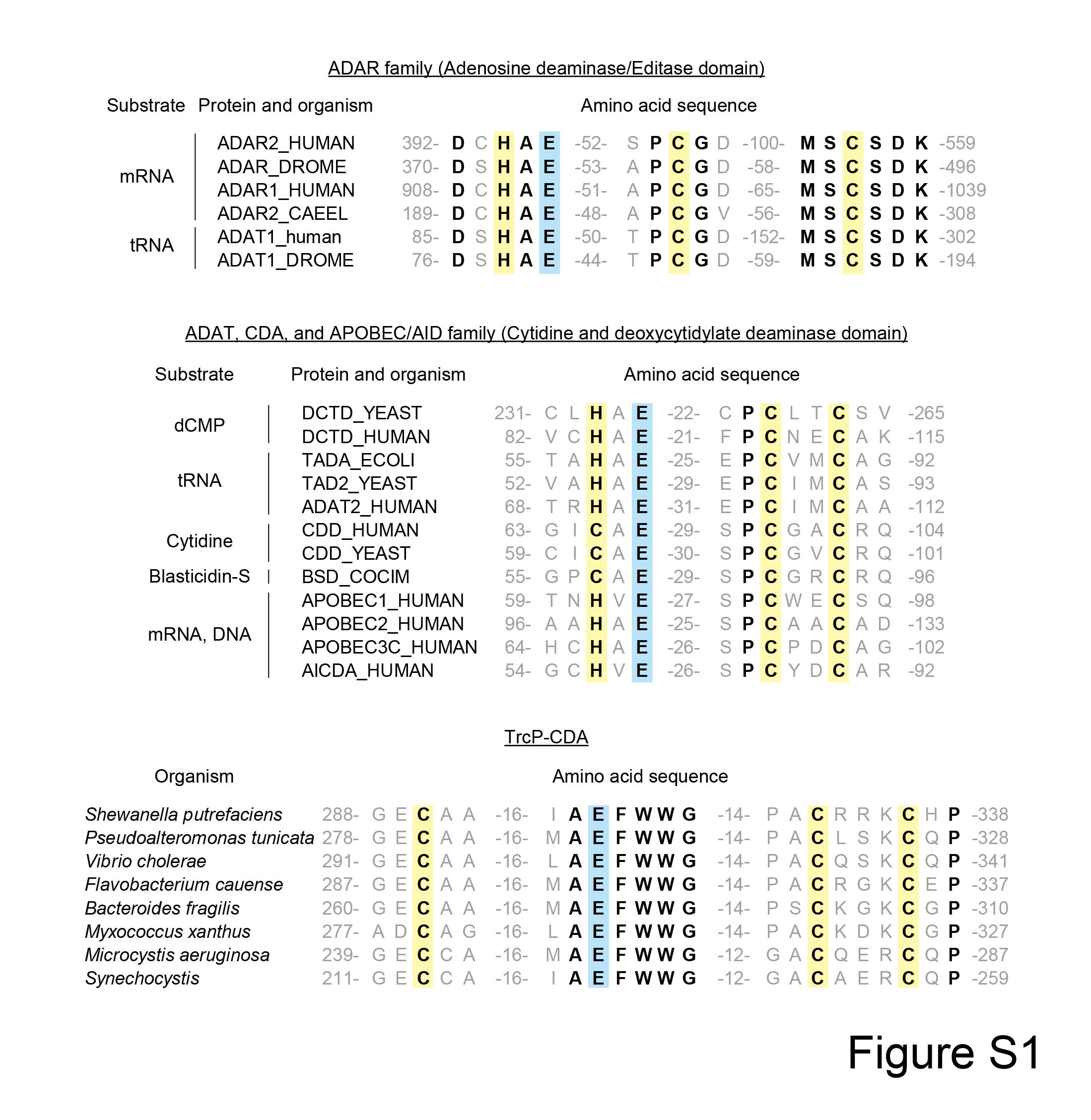

**Figure. S1 Sequence motifs in deaminases.**

Conserved histidine/cysteine clusters for coordination of zinc are highlighted in yellow. The catalytic glutamate residue is highlighted in light blue. The substrate molecules for the ADAR, ADAT, CDA, and APOBEC/AID families are shown on the left. The amino acids that are identical between all sequences are shown in bold. The numbers represent the first and last amino acid positions of the retrieved sequences and the length of the amino acids between conserved motifs.

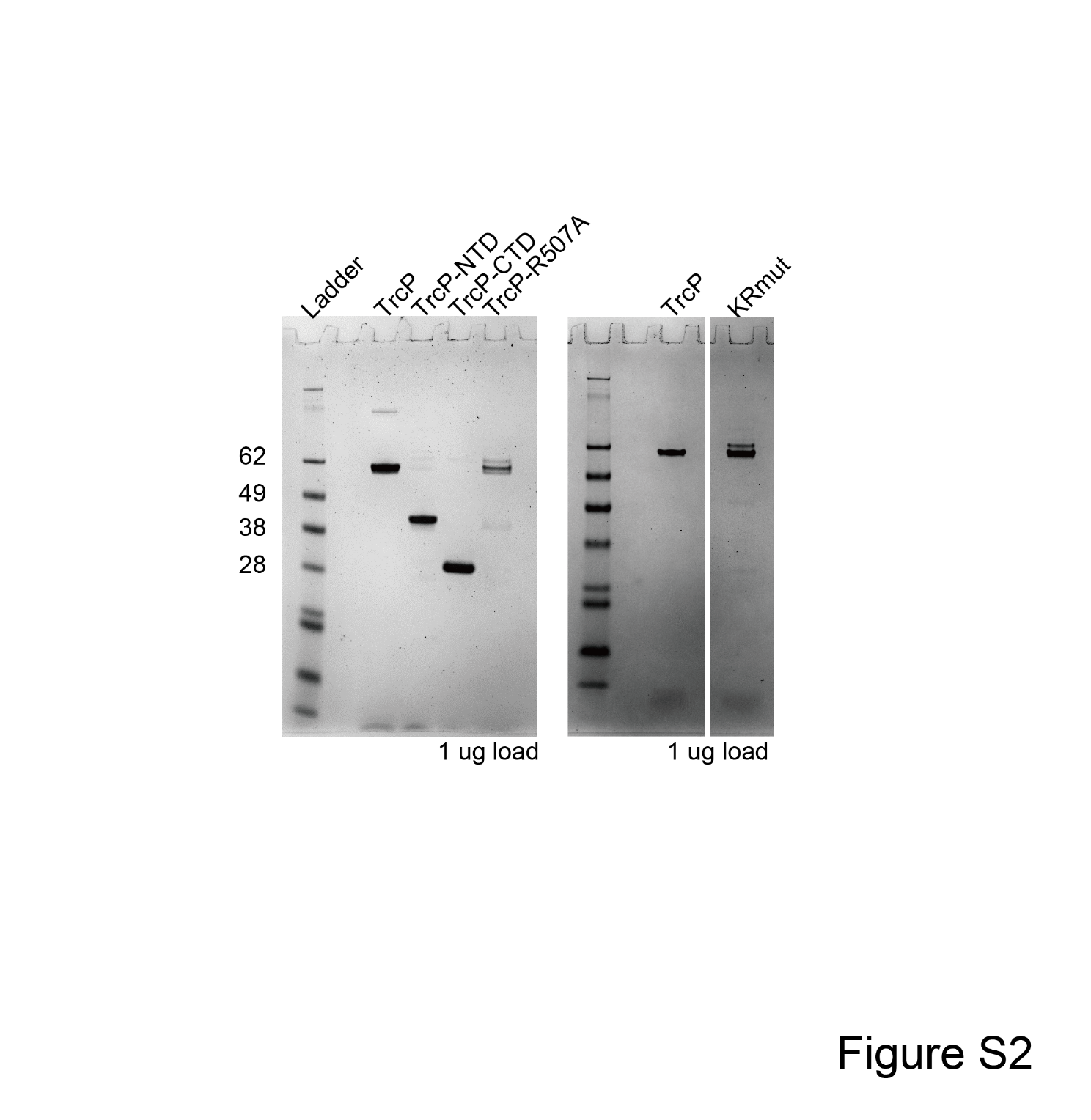

**Figure. S2 Purified recombinant proteins**

Coomassie stained gels of purified proteins; 1 μg of each protein is loaded.

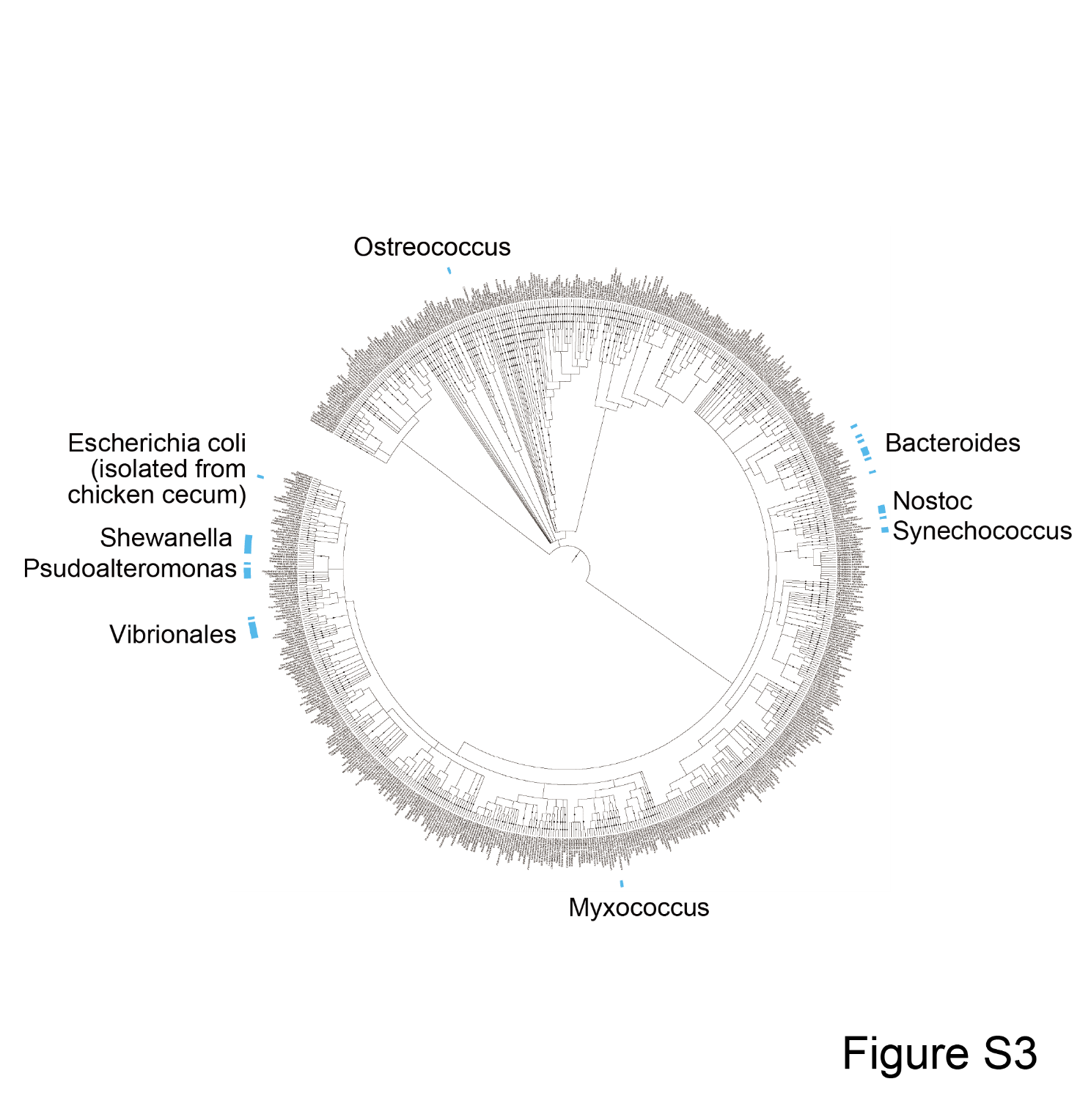

**Figure. S3 Phylogenetic tree with the distribution of TrcP homologs**

The organisms bearing TrcP homologs are indicated with light blue bars.

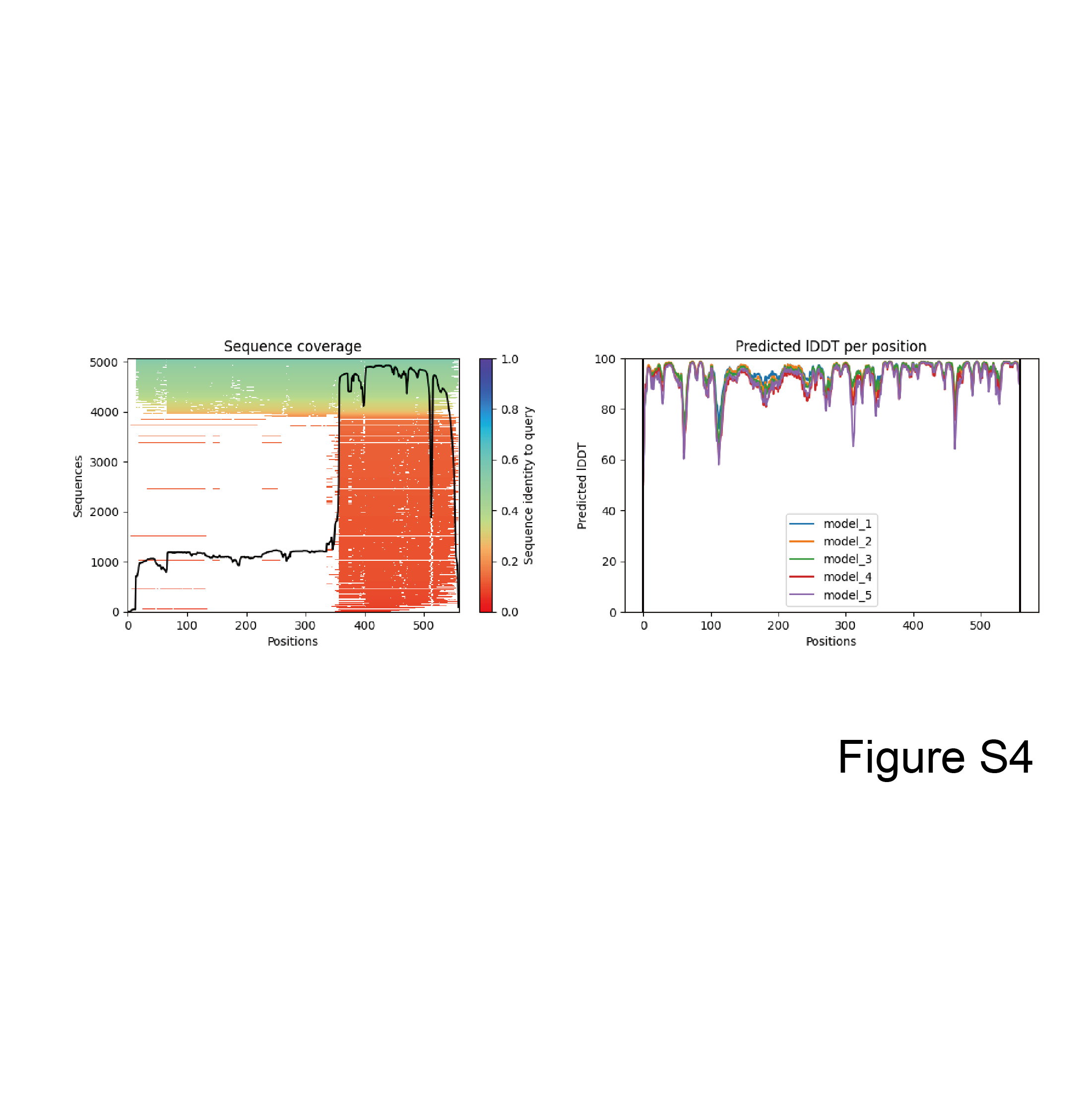

**Figure. S4 Quality scores of structural predictions by ColabFold**

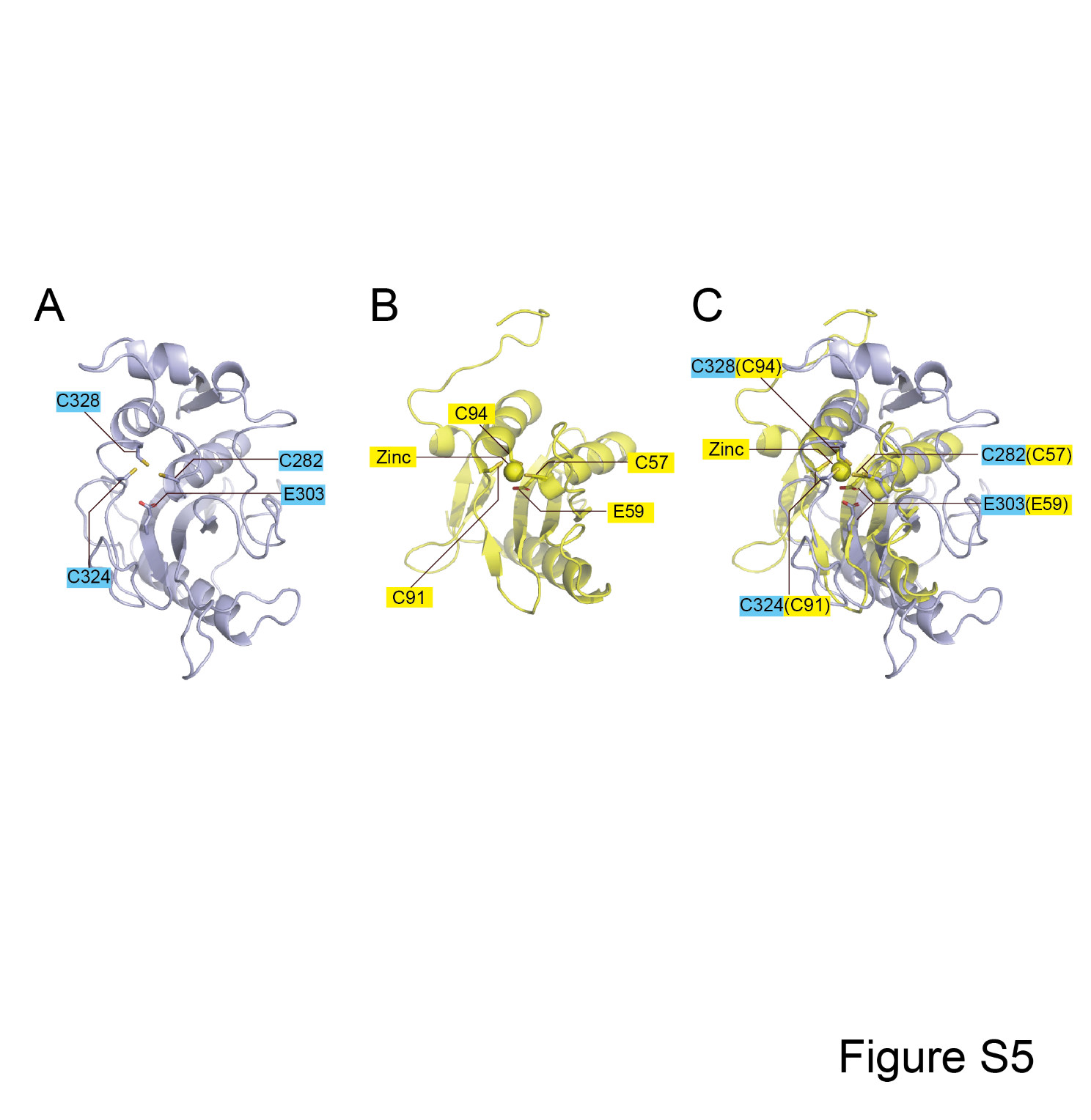

**Figure. S5 Comparison between the predicted structures of TrcP and Blasticidin-S deaminase (BSD)**

**A.** Predicted structure of the TrcP-CDA domain. The conserved cysteine and glutamate residues are indicated.

**B.** Structure of Blasticidin-S deaminase (PDB 3oj6). The cysteine cluster and conserved glutamate residue in the catalytic site and zinc ion are shown.

**C.** A structural alignment of TrcP-CDA domain and BSD. The conserved cysteines, glutamate and zinc ion are shown.

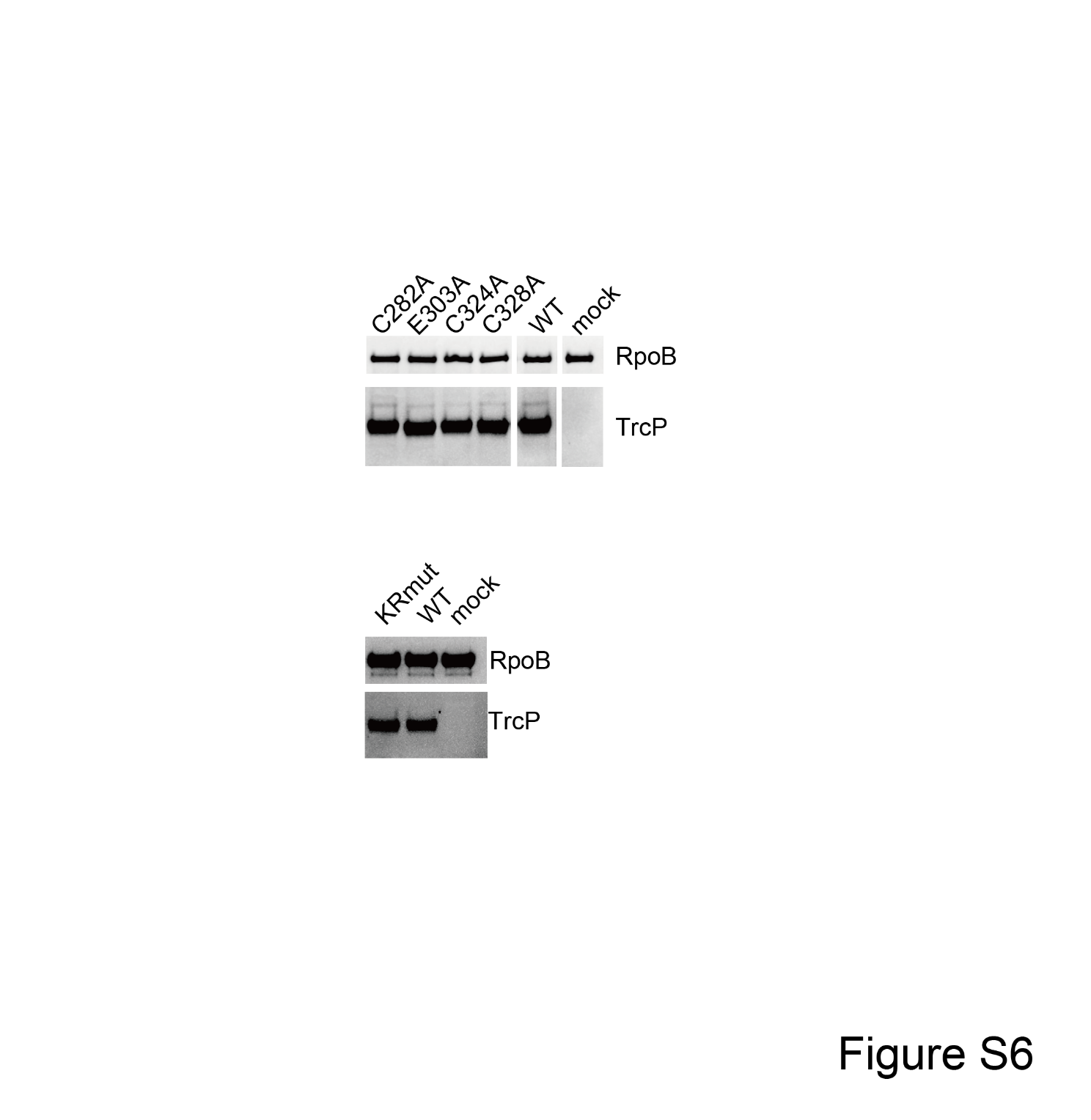

**Figure. S6 Expression levels of mutant TrcP proteins in strains used for *in vivo* complementation assays.**

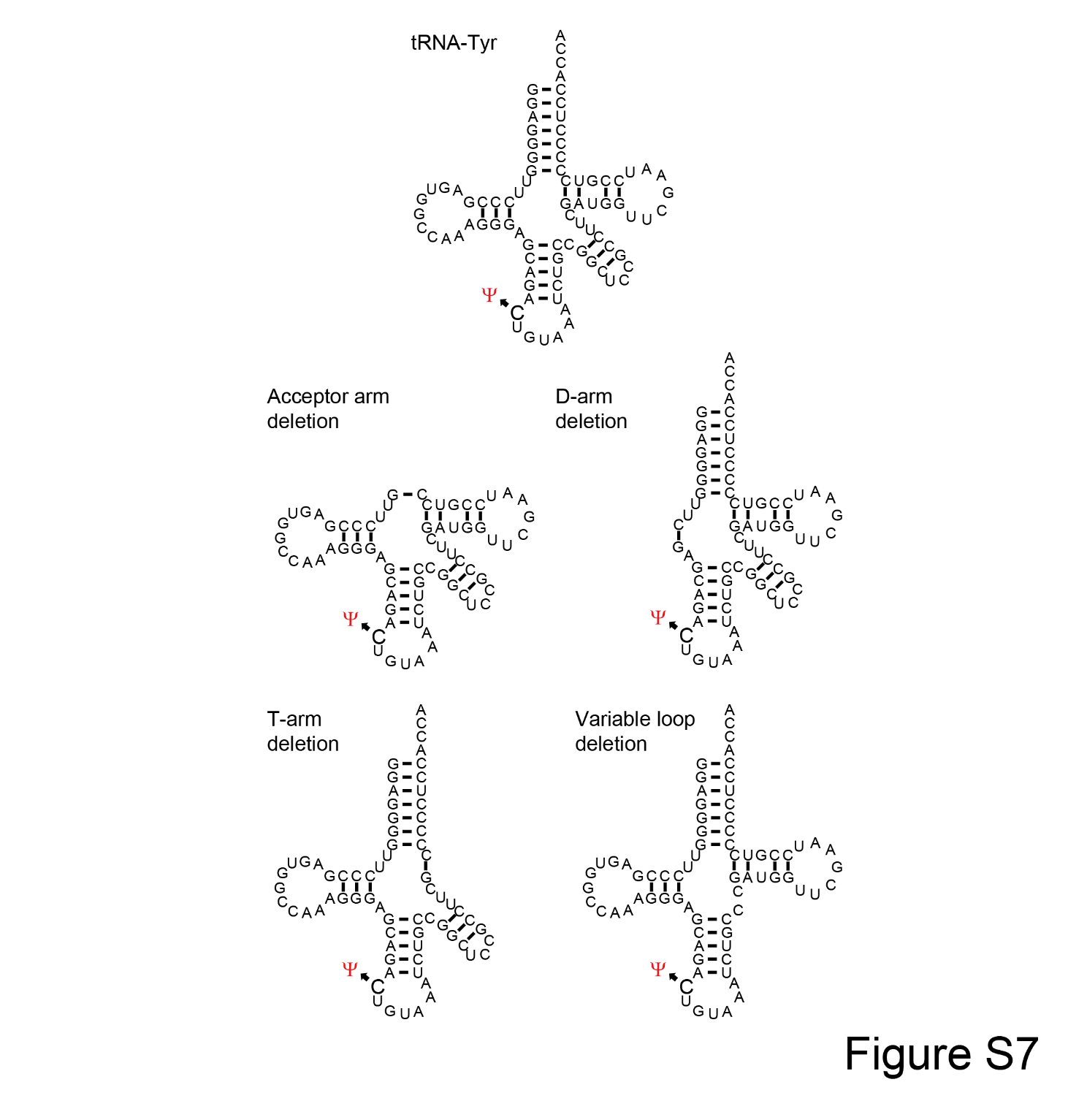

**Figure. S7 Secondary structures of tRNA-Tyr mutants tested in Figure. 4D**

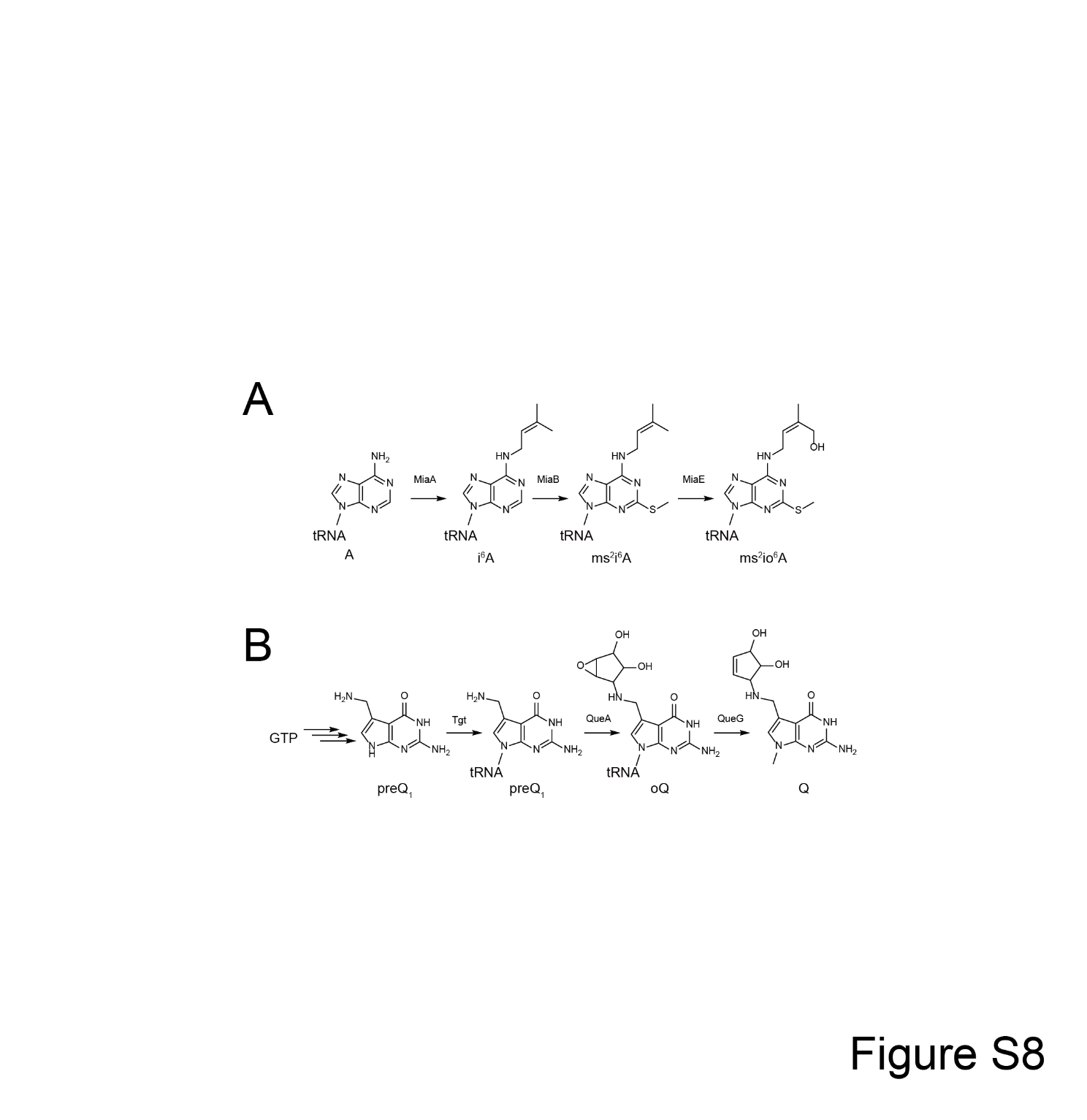

**Figure. S8 Biosynthesis pathways of ms^2^io^6^A (A) and Q (B)**

**Supplementary Data 1 Structural homology search using full-length TrcP as a query**

**Supplementary Data 2 Structural homology search using the TrcP-CDA domain as a query**

**Supplementary Data 3 Strain list**

**Supplementary Data 4 Plasmid list**

**Supplementary Data 5 Primer list**

**Supplementary Data 6 Parameters of mass spectrometry for dynamic MRM analyses**

**Supplementary Data 7 A result of Blast search using the TrcP-NTD as a query**

**Supplementary Data 8 Phylogenetic distribution of TrcP-NTD homologs**
